# Neurological Imprints of Boys’ Love Culture: Shaping Sexual Orientation Identity in the Digital Age’s Young females

**DOI:** 10.1101/2023.09.18.558088

**Authors:** Na Ao, Xiaowei Jiang, Yanan Chen, Yingying Chen, Huihui Niu, Shuoyan Hu

## Abstract

With the omnipresence of social media, Boys’ Love (BL) culture has found a burgeoning audience among young females. While this cultural phenomenon offers a platform for self-expression to some, its potential implications remain underexplored. This study investigates whether immersion in BL culture impacts the process of young females forming self-recognition and identification of sexual orientation. Employing an fNIRS-based experiment focusing on the prefrontal cortex, we compared the neurological responses of young females within and outside the BL media cocoon when exposed to sexuality-related stimuli. Our findings revealed significant neural differences in the experimental group viewing BL images compared to the control group. These results provide fresh insights into how BL content may shape our cognition and attitudes, emphasizing the need for guidance in content consumption among young females.

## Main

In the vast digital expanse of the 21st century, media has evolved beyond a passive consumable, emerging as a dominant force that molds the cognitive, emotional, and neurological pathways of its audience. Amidst this transformation, Boys’ Love (BL) culture, originating from Japan, stands out, illustrating the profound impact media can wield^1^. This niche genre, which presents romantic liaisons between male protagonists, has remarkably enamored a global female audience, particularly young females^2^. But this is more than a cultural quirk: the proliferation and nuances of BL culture compel us to reevaluate and deepen our understanding of media’s pervasive influence on societal perceptions.

Conjure a world where sexual orientation’s boundaries, traditionally delineated by biology, societal expectations, and personal experiences, could be subtly influenced by the media we voraciously consume. An fMRI study postulated that human sexual preference might be linked to neuronal responsiveness in the reward and motor systems when engaging with sexually suggestive content. Another critical research emphasized that our sexual orientations are possibly imprinted during early fetal phases, influenced by a tapestry of genetics, prenatal hormones, and unique maternal immunization processes^3^. Such revelations underscore the necessity of assessing the potential ramifications of globally accessible BL narratives on the burgeoning minds of adolescents and young females.

The digital age, marked by the ubiquity of social media, has obliterated geographical confines, ushering in a new paradigm where content is globally accessible. Consequently, even cultures once insulated from BL narratives now find themselves enmeshed within its allure. This phenomenon transcends being a mere trend; it signifies a global media consumption revolution, potentially altering the societal foundations we’ve taken for granted.

BL content’s charm is manifold^4^. By offering an alternative to the conventional heteronormative scripts and shedding light on the emotional intricacies of male characters often sidelined in mainstream narratives, it captivates and resonates with its audience, occasionally even recalibrating perceptions. Groundbreaking studies offer empirical evidence, revealing that neural responses to sexual stimuli can vary significantly based on one’s orientation, emphasizing the gender-specific nature of men’s responses compared to the more diverse responses in women^5,6^.

In this transformative journey of self-discovery and identity, young females emerges as a pivotal phase, characterized by questioning societal norms and forging personal beliefs^7^. Within this context, BL media’s ‘information cocoon’—an environment where individuals are predominantly exposed to reinforcing information—assumes profound significance. For the young female demographic engrossed in BL narratives, this suggests a milieu where male relationships are not just normalized but potentially idealized. Given this backdrop, and armed with insights about the pliability of young female neural pathways regarding sexual identity ^8,9^, and the potential neural ramifications of prolonged BL media immersion demand earnest attention.

Our study was a foray into this uncharted territory. We firstly do a survey about the attitude to boy’s love of these young female (see Supplementary Information for the complete survey description). Employing behavioral metrics coupled with fNIRS methodologies, we then probed the cognitive implications of BL culture on young females. Our comprehensive statistical analysis unveiled the attitude and gender identify changing and discernible brain response patterns between females exposed to BL content and those untouched by its influence. While the former exhibited uniform brain responses to both boy’s love and heterosexual stimuli, the latter manifested heightened brain activity for boy’s love content. This disparity in neurological reactions accentuates the profound influence consistent BL content exposure can have on the viewer’s psyche.

Navigating the digital era, characterized by media’s overarching influence on young females’ worldview, discerning the nuanced impacts of niche genres like BL becomes indispensable. Our investigation illuminates the intricate nexus between media consumption, particularly BL culture, and its profound cognitive and neurological consequences. As media narratives continue to evolve and redefine societal norms, grasping the multifaceted repercussions of niche cultural immersion is crucial. Our study accentuates the pivotal role such understanding assumes in contemporary society, offering invaluable insights for educators, policymakers, and guardians. In tandem, insights from studies, which elucidate the genetic intricacies surrounding same-sex sexual behavior, further enrich our comprehension, underscoring the evolutionary complexities and potential advantages linked to same-sex oriented behaviors^10^. As we stride forward, equipping our younger generations to critically engage with digital narratives becomes paramount, ensuring a balanced and informed perspective in this ever-evolving digital age.

## Result

### Behavior Results

#### Reaction Time

Analysis revealed a pronounced reaction time difference across image types (*F* (2,58) = 135.412, *p* < 0.001, *η_p*^^2^ = 0.824). However, distinctions between groups (fans vs. non-fans) were not statistically significant (*F* (1,29) = 1.247, *p* > 0.05, *η_p*^^2^ = 0.041), and neither was the interaction effect (*F* (2,58) = 2.319, *p* > 0.05, *η_p*^^2^ = 0.074), participants exhibited prolonged reaction times (Fig. 2B) for boy’s love images (M = 1066.987 ms, SD = 213.255,*t*=6.121,*p*<0.001) and heterosexual images (M = 1059.496 ms, SD = 213.196,*t*=5.969,*p*<0.001) compared to mask counterparts (M = 767.596 ms, SD = 139.372).

#### Accuracy

Image categorization accuracy varied significantly (*F* (2,58) = 22.622, *p* < 0.001, *η_p*^^2^ = 0.439), yet neither group nor interaction effects reached significance (*F* (1,29) = 1.921, *p* > 0.05, *η_p*^2 = 0.062; *F* (2,58) = 1.662, *p* > 0.05, *η_p*^^2^ = 0.054, respectively). Refer to Fig 2B, the accuracy in recognizing male boy’s love images (M = 0.952, SD = 0.035, *t* =-6.048, *p*<0.001) and heterosexual (M = 0.972, SD = 0.024, *t*=-4.356, *p*<0.001) less than mask images (M = 0.994, SD = 0.014). Specifically for non-fans, boy’s love images accuracy (M = 0.940, SD = 0.039) was inferior (*t*=-2.436, *p*<0.05) to heterosexual images (M = 0.970, SD = 0.024).

### fNIRS Results

#### Statistical Results of averaged hemoglobin concentration data

Concerning the Oxy-Hb responses in the channel 19 (right ventrolateral prefrontal cortex, rVLPFC), a noteworthy interaction effect was observed (*F* (2,58) = 6.649, *p* < 0.05, *η_p*^^2^ = 0.187). In contrast, the main effects for both image type (*F* (2,58) =0.503, *p* > 0.05, *η_p*^^2^ = 0.017) and participant classification (*F* (1,29) = 0.338, *p* > 0.05, *η_p*^^2^ = 0.012) did not exhibit significance. Upon exposure to boy’s love imagery, fans registered a relatively attenuated (*t*=2.598, *p*<0.05) response (M=0.160, SD = 0.061) when compared to non-fans (M=0.221, SD = 0.070), demonstrating statistical significance (*p*<0.05). Conversely, non-fans exhibited a less pronounced reaction (*t*=2.553, *p*<0.05) when presented with heterosexual imagery (M = 0.163, SD=0.049) than BL images (M =0.221, SD=0.070). These results are graphically represented in Fig 2A, B,C, and E.

## Discussion

Our study, set against the backdrop of the burgeoning popularity of Boys’ Love (BL) culture, sought to understand the neurological implications of BL media exposure on young females, we found female fans are first exposed to BL in 16±2.2 years old. As the results unraveled distinct neural patterns between the BL-exposed and unexposed groups, particularly concerning the perception of boy’s love stimuli, several discussions emerge that both validate and challenge existing paradigms.

### Impact of BL Media Cocoon on Neurological Perception

The noticeable absence of neurological differentiation between boy’s love and heterosexual stimuli within the BL-exposed group represents a significant departure from established norms. At its core, this discovery suggests that continuous immersion in BL content might potentially blur or modify traditional lines of sexuality perception. This aligns with the notion of the ‘information cocoon,’ wherein sustained exposure to a singular narrative or viewpoint can significantly shape an individual’s perception and understanding of certain topics.

The passive nature of this altered perception, predominantly an outcome of consistent media consumption, raises essential questions regarding the boundaries between media influence and inherent neural processes. The human brain, particularly during the impressionable young phase, is known for its plasticity, adaptability, and malleability^11^. The capacity of the brain to adapt to external influences has been widely documented, with environmental stimuli often playing a role in the formation of neural pathways^12^. Our findings, set against this understanding, suggest that BL content can potentially serve as one such influential external stimuli, steering neurological perceptions about romantic and sexual orientations.

### Neural Signatures of BL Perception in the Prefrontal Cortex

In our exploration of neural responses to BL media exposure, one specific area emerged with pronounced significance: the right ventrolateral prefrontal cortex (rVLPFC). Historically, the rVLPFC has been implicated in tasks involving decision making, emotion regulation, and self-recognition ^**13,14,15**^. Given these traditional functions, its heightened activity in BL-exposed individuals prompts intriguing interpretations.

One potential explanation is that this region might mediate the process of integrating unconventional romantic and sexual narratives, such as those presented in BL media, into the individual’s cognitive framework. Continuous exposure to BL content might engage the rVLPFC in a way that facilitates the accommodation and normalization of these narratives, especially given the average age of first exposure being 16±2.2 years, a time as a key period of brain rewiring^**16**^.

Moreover, the differential neural pattern observed in the rVLPFC between BL-exposed and unexposed groups underscores its potential role in shaping sexuality perception and emotion regulation for the young females^**17**^. It is conceivable that the rVLPFC, as part of a broader neural network, plays a critical role in determining how BL narratives are encoded, processed, and eventually integrated into one’s conceptual understanding of relationships^**18**^. Exposure to BL content may initially elicit positive emotions towards the homosexual group among fans, subsequently resulting in a decrease in activity within the right ventrolateral prefrontal cortex (rVLPFC) when viewing BL stimuli^**17**^. Furthermore, prolonged exposure to an BL media cocoon characterized by excessively idealized depictions of same-sex romantic relationships may potentially lead to a shift in the heterosexual orientation identification among fan groups.

To summarize, our results highlight the central role of the rVLPFC in modulating the neural perception of BL stimuli, suggesting a deep intertwining of media exposure, cognitive processing, and neurological responses^**19**^. The interplay between these factors, especially in the backdrop of young, underscores the profound influence of media on shaping and redefining our neurological and perceptual landscape^**20**^.

### Attitudinal Shift and its Societal Implications

Beyond the neurological differences, the broader societal implications of our findings warrant discussion. The marked neural differentiation between the two groups when viewing boy’s love images underscores the power of media in shaping or reinforcing attitudes. One could argue that the BL media cocoon, in normalizing homosexual relationships, can act as a counterbalance to traditional or societal biases against homosexuality^21^.

This aligns with the broader discourse on media representation and its role in challenging stereotypes. Historically, marginalized groups have often found solace and representation in niche media. BL, by its very nature, offers a space where homosexual relationships are celebrated, thereby possibly playing a role in destigmatizing and normalizing such relationships for its audience.

However, it’s also worth noting that BL narratives are fictional and often idealized. The risk lies in young females developing a romanticized or skewed perception of real-world homosexual relationships and even denying one’s own female identity based on these idealized depictions. While the destigmatization of homosexual relationships is undoubtedly a positive outcome, it’s essential to differentiate between representation and reality.

### Ethical and Educational Implications

Our findings, particularly concerning the potentially malleable nature of neurological perceptions during young, also introduce ethical considerations. The transformative potential of media content, especially when consumed uncritically, underscores the importance of media literacy education. As revealed by our study in the supplementary materials, contrary to expectations, female fans did not exhibit higher educational attainment compared to their peers in cases where parents had higher levels of education. By equipping young individuals with the skills to critically analyze and evaluate media content, we can ensure a more balanced and informed consumption pattern.

Additionally, it’s worth pondering upon the accessibility and regulation of such content. The recommendation to potentially regulate content accessibility based on age, as suggested in the conclusion, can be seen both as a protective measure and a restriction on creative freedom. Striking a balance between safeguarding young minds and encouraging free expression remains a challenge that requires collective societal deliberation.

### Future Research Directions

While our study has unveiled specific neurological patterns associated with BL media consumption, there remains a vast terrain to explore. Future research could delve into longitudinal studies to understand the long-term effects of such exposure. Additionally, exploring diverse cultural contexts can provide a more comprehensive understanding of the global impact of BL culture. Comparing the neural effects of different genres of media content, beyond just BL, can also offer insights into the broader discourse of media influence.

### Concluding Thoughts

In an era where media has become an inseparable facet of daily life, understanding its multifaceted impact is not just an academic endeavor but a societal imperative. Our study, rooted in the intersection of media culture and neuroscience, serves as a step towards deciphering the intricate web of influences that shape our perceptions, beliefs, and identities. As the landscape of media consumption continues to evolve, so must our efforts to comprehend, navigate, and guide it.

## Method

### Participants and Procedure

#### Recruitment and Sample

A total of 151 (fans = 80, nonfans = 71) and 40 (fans = 20, nonfans = 20) female undergraduates from Henan University were enlisted for the Image-Rating and Image-Classification tasks, respectively. 12 participants from the Image-Rating task and 19 from the Image-Classification task was excluded due to protocol non-adherence or excessive head movement. This left 139 participants for Image-Rating (73 fans and 66 non-fans) and 31 for Image-Classification (17 fans and 14 non-fans). Participants received 30RMB for their contribution.

#### Procedure

The Image-Rating task had participants rate 353 images. 120 images were selected based on these ratings for the subsequent Image-Classification task. Before experiments, participants provided written consent, filled a demographic questionnaire, and were screened for exclusions such as neurological disorders and recent drug use. The study was green-lit by Henan University’s Ethics Committee.

#### Stimuli

Image Rating task: In this task, 353 images, both boy’s love and heterosexual, were sourced from public databases. Participants rated each image’s arousal and valence on a 1-9 Likert scale. Each trial initiated with a 2000 ms central black fixation cross, followed by a randomly presented image awaiting ratings, The flowchart is shown in Fig. 1A Image Selection: To avoid confounding valence differences with arousal, boy’s love images and heterosexual couple images were matched on arousal(*t* (118) = 0.265, *p* > 0.05) and valence (*t* (118) = 1.042, *p* > 0.05) and using normative ratings from fans (arousal: *t* (118) = 0.043, *p* > 0.05, valence: *t* (118) = 0.407, *p* > 0.05) and nonfans (arousal: *t* (118) = 1.302, *p* > 0.05, valence: *t* (118) = 0.100, *p* > 0.05). We then scrambled the images by dividing the boy’s love images into segments (each fragment was 20 Pixels×20 Pixels) and then randomly recombined the segments in MATLAB, illustrated in Fig. 1B. The mask images 1) removed meaningful content; 2) changed the position of the spatial frequency components; and 3) maintained the same lower-level visual properties as the original image. All images were matched on average luminance using the SHINE MATLAB toolbox ^22&23^ .

**Fig. 1.**
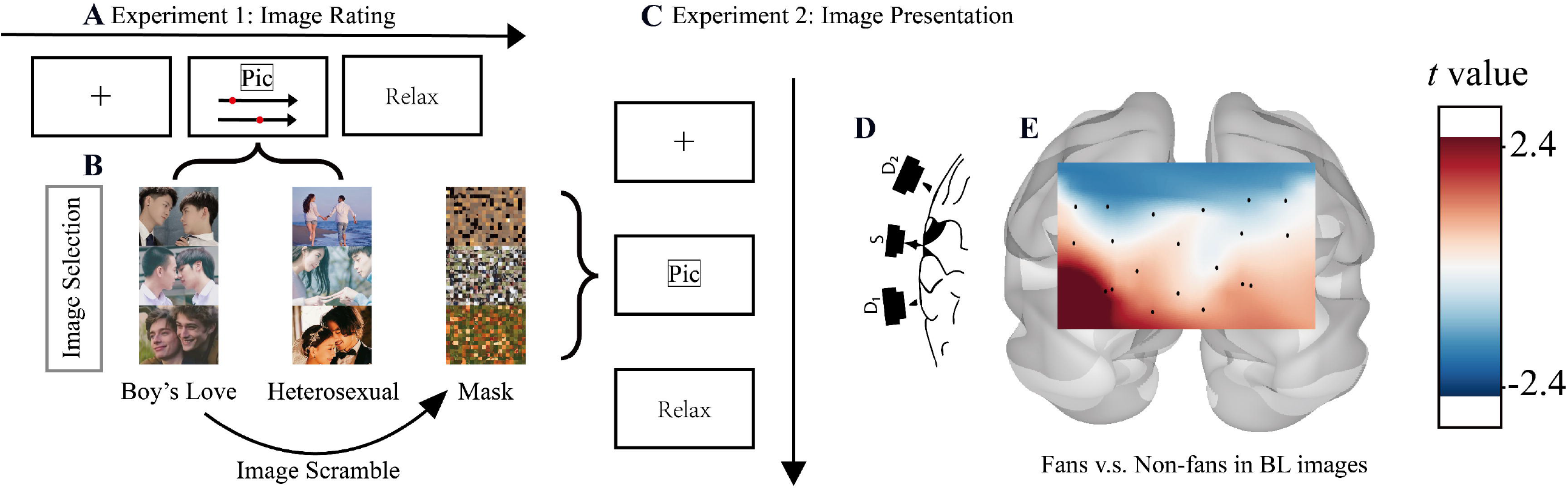
Experiments Flow Chart. **A**, The experiment 1 (imaging rating) begins by asking the participants to rating 2 features (arousal and pleasure) for each image. **B**, Experiment 2 (image classification) after balancedly selecting images from the experiment 1, participants classify the presented pictures and mask images are generated by scarabling the Boys’ Love (BL) images. **C**, The fNIRS experiments begins by persenting the selected image to the participants and record their behavioral and neural responses. **D**, the schematic of how to record the signals from the fNIRS channel by sending the light from sources to detectors. **E**, The *t*-value topomap of comparing the fans and non-fans while persenting the BL images.

**Fig. 2.**
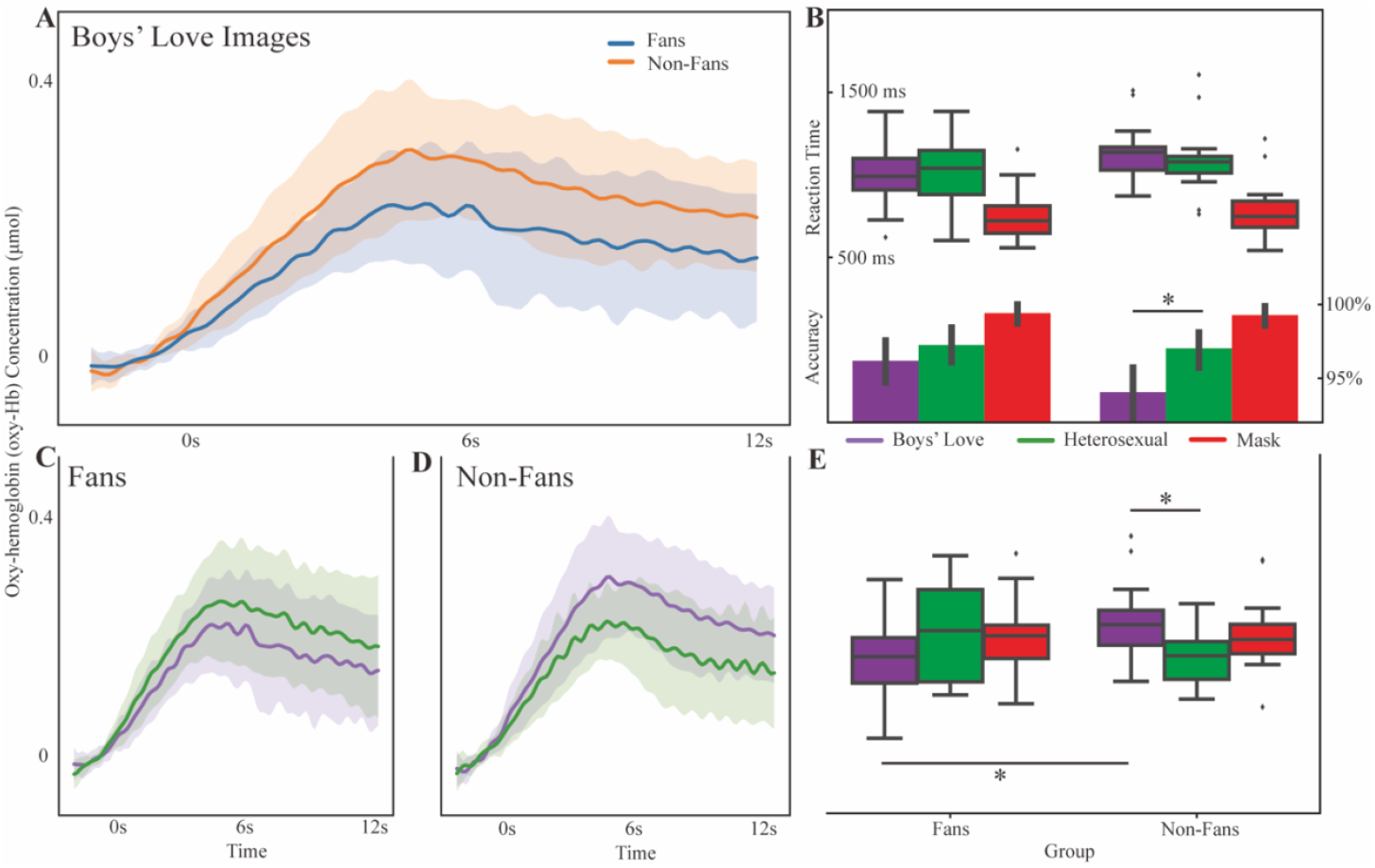
The behavioral and main neural results. **A**, The Oxy-hemoglobin (oxy-Hb) concentration (μmol) signal of comparing the fans and non-fans while persenting the BL images in channel 19, the shallow area shows the standard error bars. **B**, The behavioral results (reaction time and accuracy) of two groups (left: fans, right: non-fans); *: p < 0.05. **C**, The comparation of the oxy-Hb concentration (μmol) signal while persenting Boys’ Love (BL) and heterosexual images to fans. **D**, The comparation of the oxy-Hb concentration (μmol) signal while persenting Boys’ Love (BL) and heterosexual images to non-fans. **E**, The averaged oxy-Hb concentration (μmol) responses for 3 (images) × 2(groups) stimuli.

Image-Classification task: Post 5-minute rest with fNIRS headcap, participants embarked on the Image-Classification task. They navigated 180 trials, discerning between images of boy’s love, heterosexual couples, or mask images (60 each type). Task architecture (Fig. 1C) comprised: a 2000 ms fixation cross, 4000 ms target (400 x 300 pixels), and 8000 ms blank post-response. Reaction time, accuracy, and fNIRS metrics were documented.

### fNIRS Data Collection

#### Acquisition

We employed a 20-channel fNIRS system (NIRX Scout 32x32, USA) at 7.8125 Hz sampling rate to log oxy-Hb, deoxy-Hb, and total-Hb absorptivity. This configuration, encompassing 15 probes (8 sources x 7 detectors; the schematic was shown in Supplementary materials Fig. 1), with an inter-probe distance of 30 mm, was concentrated over the prefrontal cortex, giving 20 channels. Dual wavelengths (785 nm, 830 nm) minimized crosstalk (Supplementary materials Fig. 1). Probe alignment followed the International 10-20 system, centering at FPz. Channel topography on standard cortex was visualized using NirsScan (Fig 1E).

#### Behavioral Analysis

Image classification accuracy (%) and reaction time (ms) were evaluated using a 3x2 mixed ANOVA in SPSS 25(2017). The within-subject factor was image type with three levels: boy’s love, heterosexual couple, and mask images. The between-subject factor was participant type: erotica fans versus nonfans. Post-hoc analyses employed t-test, False positive correction by Benjamini-Hochberg.

### fNIRS Analysis

#### Preprocessing

Using MNE 0.22^24^ on Python 3.10.2, raw fNIRS light intensity data were transposed to optical density, then, the scalp coupling index (sci) was used to evaluate the channel quality of the coupling between the opcodes and the scalp^25^. If the sci of the channel was less than 0.5, the channel was rejected and interpolated. Temporal derivative distribution repair (TDDR)^26^ addressed baseline shifts and spike artifacts. Conversion to hemoglobin concentration employed the Beer-Lambert law, a band-pass filter (0.005 Hz -0.2 Hz) minimized drifts and cardiac interference, excluding anomalies exceeding a threshold of 100 mmol. Hemoglobin data were segmented from -2 s to 12 s, baselined, and then averaged within conditions.

## Statistical Analysis

Averaged hemoglobin concentrations were explored channel-wise. Similar to behavior analysis, a 3x2 mixed ANOVA and T-tests discerned main effects. Subsequent sample analysis compared hemoglobin averages across main effect levels.

## Supporting information

None

## Data Availability Statement

Because personal privacy is involved and some of the subjects do not want their options to be actively disclosed. All data, scripts, and images in the present study are available to require.

## Acknowledgments

We thank all participants in the present study, all students who assisted with data collection. This research was supported by Institute of Psychology, CAS (No. GJ202007), the Humanities and Social Sciences project of Ministry of Education (22YJCZH021), and the Education and teaching reform project of Psychology Education commission, Ministry of Education (20222005).

## Author Contribution

Na Ao and Xiaowei Jiang contributed equally to this work and share first authorship. Yingying Chen, Huihui Niu, and Shuoyan Hu contributed equally to this study. Yanan Chen supervised. Conceptualization, Na Ao, Xiaowei Jiang, and Yanan Chen; Data curation, Na Ao, Yingying Chen, Huihui Niu and Shuoyan Hu; Formal analysis, Xiaowei Jiang and Na Ao; Investigation, Na Ao; Methodology, Xiaowei Jiang; Project administration, Yanan Chen; Validation, Na Ao; Visualization, Yingying Chen, Huihui Niu, Shuoyan Hu; Writing – original draft, Na Ao and Xiaowei Jiang; Writing – review & editing, Yanan Chen and Xiaowei Jiang.

## Materials & Correspondence

Yanan Chen correspondence and material requests should be addressed.

