## Supplementary material for "Neurological Imprints of Boys’ Love Culture: Shaping Sexual Orientation Identity in the Digital Age’s Young females": None

### Supplementary Information

a) The Boys' Love survey

#### Introduction

The cultural landscape has undergone profound transformations with the advent of new media genres, spawning subcultures that indelibly influence individuals' perceptions, attitudes, and beliefs.[1] Among these, the Boys' Love (BL) culture, which originated from Japan and pivots around romantic narratives between male characters, has gained conspicuous traction[2]. By pushing boundaries and challenging societal norms, BL presents a unique lens through which broader debates around gender roles, sexuality, and societal acceptance are refracted. Given any media engagement invariably imprints on our worldview, it is essential to ask: How does immersion in BL culture alter one's perspectives?

Despite BL culture's soaring global popularity[3] and its probable consequential influence, a paucity of empirical research exists on the demography and attitudes of its enthusiasts. Several poignant questions arise: Do BL fans exhibit differences in sexual orientation, parental education, and family economic status compared to non-fans? Do they hold distinctive views on homosexuality? Can BL culture's allure be mapped to particular demographic strata or is its resonance more ubiquitous?

In an endeavor to bridge this knowledge chasm, our study embarks on a comprehensive exploration of the BL cultural intersection with individual demographics and attitudes. Through a systematically curated survey, we probe the potential dichotomies between BL aficionados and non-enthusiasts, especially concerning sexual orientation, attitude on homosexuality, demographic variables like parental education level, family economic status, gender identify and so on. The overarching aim is to discern whether engagement with BL culture both shapes and mirrors the intricate tapestry of its audience's identities and convictions.

#### Method

##### Participant and procedure

Participants and Methods:

We conducted an online survey that successfully engaged a total of 1301 female participants, utilizing the online survey platform (https://www.wjx.cn/). Before beginning the survey, all participants were provided with a clear articulation of the research purpose and were required to sign an informed consent form, ensuring they comprehended the scope and intention of the study.

Upon completion of the survey, participants were remunerated with an amount ranging from 5 to 15 yuan as a token of appreciation for their time and participation. However, not all submissions could be considered valid for analysis. From the initial pool, 158 participants were excluded. The exclusions were due to 80 participants failing the polygraph section of the survey, while 78 were eliminated due to their response times being below 1.5 standard deviations of the average answer time[4]. This rigorous filtering process left us with 1143 valid entries for our analysis.

The average age of the final participant group was 21.54 years with a standard deviation of 4.23 years. Detailed demographic variables and other pertinent information related to these participants, which constitute our result (Table 1).

#### Material

The survey instrument deployed in this study is a multi-faceted questionnaire composed of three distinct sections. To safeguard the integrity of the results and preempt potential deceit or insincerity in answers, polygraph tests were interspersed randomly throughout these sections.

Demographic Variables Questionnaire: This section solicited fundamental demographic details from the participants, encompassing aspects such as gender, age, home environment, and sexuality. Importantly, it also gauged participants' attitudes toward BL fans.

Kinsey Scale[5]: The Kinsey Scale serves as a seminal instrument in the understanding of human sexuality. Devised to capture the spectrum of sexual orientations, it scores participants on a range from 0 to 6. In this continuum, a score of 0 signifies an exclusive heterosexual orientation, a score of 6 denotes an exclusive homosexual orientation, and a mid-point score of 3 represents a balanced orientation, i.e., equally heterosexual and homosexual. For the purposes of our study, an additional categorization, symbolized as "X", was incorporated to represent individuals identifying with asexuality.

Attitudes toward Lesbians and Gay Men in Chinese (CATLG): Developed by Yong et al[6], the CATLG is a self-report questionnaire tailored specifically for the Chinese cultural milieu. Comprising 20 items, it delineates attitudes into two primary dimensions:

Attitudes toward Lesbians (ATL): This segment contains 10 items that evaluate respondents' views regarding lesbians.

Attitudes toward Gay Men (ATG): This component, too, is constituted by 10 items, but focuses on perspectives related to gay men.

For both ATL and ATG, participants express their agreement or disagreement with statements using a 5-point Likert scale, ranging from 1 (Strongly Disagree) to 5 (Strongly Agree). The robustness and reliability of CATLG is underscored by its high internal consistency (Cronbach's $\alpha = 0.898$) and a test-retest coefficient of 0.959.

#### Result

The results reveal discernible differences between fans and non-fans of the Boys' Love (BL) culture across various parameters (Table 1). Specifically, BL fans, whether before the age of 18 or in their current place of residence, are more developed, and their parents' educational levels generally reporting higher levels. While, the fans don't show higher educational level. Additionally, BL fans exhibit a trend of 'late marriage and low birth rates' and, at the same time, are more dissatisfied with their gender as female. In terms of sexual orientation and experiences, BL fans and non-fans show comparable behaviors, but there are significant variations in their self-reported sexual orientations. Lastly, attitudes towards homosexuality, both male and female, are markedly more positive among BL fans. Interpersonal preferences also differ, with non-fans showing a reduced inclination to befriend BL fans compared to vice versa.

#### Discussion

The burgeoning prevalence of the Boys' Love (BL) culture has catalyzed an imperative inquiry into the potential attitudinal and demographic differences between its aficionados and those relatively unacquainted with it. As evidenced by the presented results, this dichotomy provides a rich tapestry of insights that can inform our understanding of the sociocultural ramifications of such subcultures.

**Demographics and Family Background:** Urban environments, often hubs of progressive ideas, correlate with a higher inclination towards the BL culture. The fact that BL fans predominantly hail from urban settings and from homes where higher education is valued underscores the role of both environment and upbringing in shaping openness to diverse narratives. This suggests a need for a wider spread of such tolerant environments to nurture acceptance and understanding of non-traditional media genres.

**Perspectives on Relationships and Sexuality:** BL narratives, characterized by their deviation from societal norms, appear to be influencing or at least resonating with fans' progressive views on marriage, child-rearing, and gender fluidity. The parallel between the liberal themes in BL content and fans' openness to unconventional relationship dynamics is undeniable. As this genre gains traction, educators might consider integrating discussions about such media in curricula, promoting critical engagement and reflection among students.

**Sexual Orientation and Experience:** While sexual experiences of both groups were largely analogous, a nuanced difference in self-identification of sexual orientation was observed among BL fans. This trend beckons an exploration into the intricate dance between media consumption and personal sexual identification. It raises the question: does BL culture attract those with broader sexual self-perceptions, or does it play a part in widening their understanding of sexuality?

**Attitudes Towards Homosexuality:** The starkly positive attitude of BL fans towards homosexuality is testament to the power of narrative immersion. By predominantly portraying homosexual relationships in a positive light, BL narratives could be fostering empathetic and supportive worldviews among their readership. This highlights the imperative for media creators to approach content creation with responsibility, recognizing the profound implications their work can have on societal attitudes.

**Interpersonal Interactions:** The palpable hesitation among non-fans to associate with BL aficionados hints at a broader societal bias. This emphasizes the need for initiatives aimed at dispelling misconceptions and promoting inclusivity.

In conclusion, the burgeoning BL subculture stands as a beacon in the digital landscape, potentially steering the evolution of young minds in their perceptions of sexuality and relationships. The interplay between BL media and its consumers sheds light on the intricate ways in which modern media shapes, reflects, or even amplifies latent beliefs. It is an urgent call to educators, policymakers, and media professionals to guide the digital-savvy youth, ensuring they approach content with a balanced and critical lens, appreciating the profound ways it might shape their worldviews.


**Table 1.** The result of the questionnaires

| **Item** | Nan-fans (N=355) | Fans (N=788) | t-value |
|---|---|---|---|
| [a] Current Residence | 1.515 (0.782) | 1.400 (0.681) | -2.536* |
| [a] Residence before the age of 18 | 2.068 (0.848) | 1.832 (0.830) | -4.403*** |
| [a] Principal guardian before18 | 1.073 (0.272) | 1.065 (0.280) | -0.480 |
| Father's level of education | 3.724 (1.682) | 4.321 (1.820) | 5.254*** |
| Mother's level of education | 3.400 (1.641) | 3.942 (1.837) | 4.775*** |
| Mine level of education | 6.896 (0.839) | 6.860 (0.760) | -0.705 |
| Family socioeconomic status | 4.727 (1.530) | 4.841 (1.436) | 1.223 |
| The prospective age of marriage | 3.977 (1.494) | 4.510 (1.858) | 4.754*** |
| The prospective number of children | 3.118 (2.210) | 3.198 (2.536) | 0.511 |
| Do you agree that females should change themselves to male | 3.856 (1.242) | 4.043 (1.208) | 2.398* |
| Do you agree that female's feel is essential in the sex activity | 4.397 (0.855) | 4.654 (0.601) | 5.812*** |
| [a] Have you ever had sex with someone of the same sex | 1.961 (0.915) | 1.937 (0.244) | -1.635 |

| | | | |
|---|---|---|---|
| [a] Have you ever had sex with someone of the different sex | 1.777 (0.417) | 1.792 (0.406) | 0.551 |
| The prospective number of sexual partners | 2.034 (0.536) | 2.109 (0.609) | 1.813 |
| The prospective number of partners | 1.980 (0.305) | 1.989 (0.408) | 0.343 |
| Your psychological gender is | 6.146 (0.128) | 5.685 (1.250) | -5.946*** |
| Your sexual orientation is | 21.259 (3.785) | 20.668 (3.921) | -2.386* |
| Are you willing to change your gender | 1.597 (1.585) | 2.260 (1.693) | 6.247*** |
| You would like to change your gender if possible | 1.980 (1.151) | 2.524 (1.294) | 6.845*** |
| Do you want to befriend the fans | 3.586 (0.963) | 4.411 (0.697) | 16.356*** |
| Do you want to be friend with the fans | 2.882 (1.046) | 3.827 (1.038) | 14.218*** |
| Attitude toward female homosexual | 38.696 (8.828) | 44.712 (5.690) | 14.129*** |
| Attitude toward male homosexual | 37.141 (8.640) | 44.853 (6.128) | 17.224*** |
| Attitude toward homosexual | 75.837 (16.518) | 89.565 (11.170) | 16.438*** |

Note: the abbreviation of Attitudes toward Lesbians and Gay Men in Chinese Scale Emotional Creativity Inventory Scale are CATLG and ECI. a mean class variable.

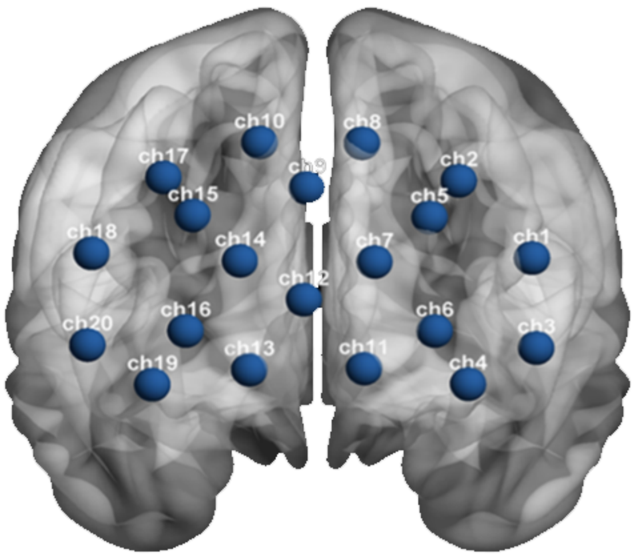


b) Fig 1. Channel location. The images illustrate the location of each channel in the prefrontal cortex.
